# A single nuclei transcriptomic analysis of the Atlantic salmon gill through smoltification and seawater transfer

**DOI:** 10.1101/2020.09.03.281337

**Authors:** Alexander C. West, Yasutaka Mizoro, Shona H. Wood, Louise M. Ince, Marianne Iversen, Even H. Jørgensen, Torfinn Nome, Simen Rød Sandve, Andrew S. I. Loudon, David G. Hazlerigg

## Abstract

Anadromous salmonids begin life adapted to the freshwater environments of their natal streams before a developmental transition, known as smoltification, transforms them into marine-adapted fish. In the wild, the extending photoperiods of spring stimulates smoltification, typified by radical reprogramming of the gill from an ion-absorbing organ to ion-excreting organ. Prior work has highlighted the role of specialized “mitochondrion-rich” cells in delivering this phenotype. However, transcriptomic studies identify thousands of smoltification-driven differentially regulated genes, indicating that smoltification causes a multifaceted, multicellular change; but direct evidence of this is lacking.

Here, we use single-nuclei RNAseq to characterize the Atlantic salmon gill during smoltification and seawater transfer. We identify 20 distinct clusters of nuclei, including known, but also novel gill cell types. These data allow us to isolate cluster-specific, smoltification-induced changes in gene expression. We also show how cellular make-up of the gill changes through smoltification. As expected, we noted an increase in the proportion of seawater mitochondrion-rich cells, however, we also identify a reduction of several immune-related cells. Overall, our results provide unrivaled detail of the cellular complexity in the gill and suggest that smoltification triggers unexpected immune reprogramming directly preceding seawater entry.

## Introduction

The Atlantic salmon migrates between fresh and seawater environments ^1^. Atlantic salmon eggs hatch in freshwater streams where they develop for 1-4 years. On reaching a critical size threshold, young “parr” animals are sensitized by several weeks of winter photoperiod (day-lengths), after which long, summer-like photoperiods stimulates the parr to transform into a marine-adapted “smolt” fish ^2^. This process, known as smoltification, drives divergent expression of endocrine factors that collectively deliver phenotypic remodeling, of length, weight, silvering, and in particular: gill physiology ^1^.

The salmonid gill is a complex multifunctional organ, essential for gas exchange, nitrogenous waste excretion, pH balance and osmoregulation ^3^. It is also a major mucosal immune barrier harboring a dedicated lymphoid tissue ^4^. Structurally, the gills are arranged in symmetrical arches, each of which are populated by numerous filament structures, which are themselves densely flanked with lamellae. The gill is composed of seven major cell types ^5^. Pavement cells (PVCs) have an enlarged surface area on the apical membrane, and form the majority of the epithelium ^6^. Pillar cells (PCs), which are structural cells, define the blood spaces within the lamellae ^7^. Goblet cells (GCs) reside in the filament epithelium and excrete mucus ^8^. Non-differentiated progenitor cells (NDCs) colonize basal and intermediate layers of the gill epithelium ^9^. Chemosensory neuroepithelial cells (NECs) lie along the length of the efferent edge of the gills and are innervated by the central nervous system ^10^. Mitochondrion-rich cells (MRCs) and their adjacent accessory cells (ACs), finally, are located at the trough between two lamellae where they abundantly express the channels and pumps required to maintain the osmotic gradients between blood plasma and both fresh-and seawater ^11&13^.

Under freshwater, Na^+^ ions are directly or indirectly exchanged for protons across the apical membrane then transported into the blood *via* the sodium potassium ATPase (NKA) on the basolateral membrane ^14–16^. Cl^−^ ions, meanwhile, are exchanged or channeled across the apical membrane then enter the blood through an undefined channel ^17&20^. Under saltwater, basolateral NKA generates a chemical and electrical gradient, motivating both loss of Cl^−^ ions *via* apical CFTR channels and paracellular escape of Na^+^ ions ^13,21^ (reviewed in ^22^).

Smoltification converts the Atlantic salmon gill from a freshwater-adapted organ to a seawater-adapted organ. Rising cortisol and growth hormone along with falling prolactin propels smoltification. This change in endocrinology coincides with a switch in anatomical and molecular phenotypes of MRCs and ACs, cell types which to date comprise the major focus of smoltification of gill physiology ^1,12,23^. Smoltification is, however, a complex developmental transition and the smolt gill phenotype likely extends far beyond changing MRC and AC cell phenotype. Therefore, to realize the complexities of smoltification-driven changes in gill physiology we adopt a new strategy, single-nuclei RNAseq, which provides transcriptional responses to smoltification and seawater transfer at individual nuclei-level resolution.

## Results and Discussion

### A single-nuclei survey of Atlantic salmon gill cells

We profiled 18,844 individual nuclei from eight Atlantic salmon gill samples from four distinct physiological states (Figure 1A). To define shared correlation structure across datasets we pooled replicate samples and integrated all four states using diagonalized canonical correlation analysis followed by L2 normalization. We next identified pairs of mutual nearest neighbors (MNNs) to identify anchors: cells that represent shared biological states across datasets. Anchors were then used to calculate “correction” vectors allowing all fours states to be jointly analyzed as an integrated reference ^24^. Unsupervised graph clustering partitioned the nuclei into 20 clusters, which we visualized using a uniform manifold approximation and projection (UMAP) dimension reduction technique (Figure 1B).

**Figure 1.**
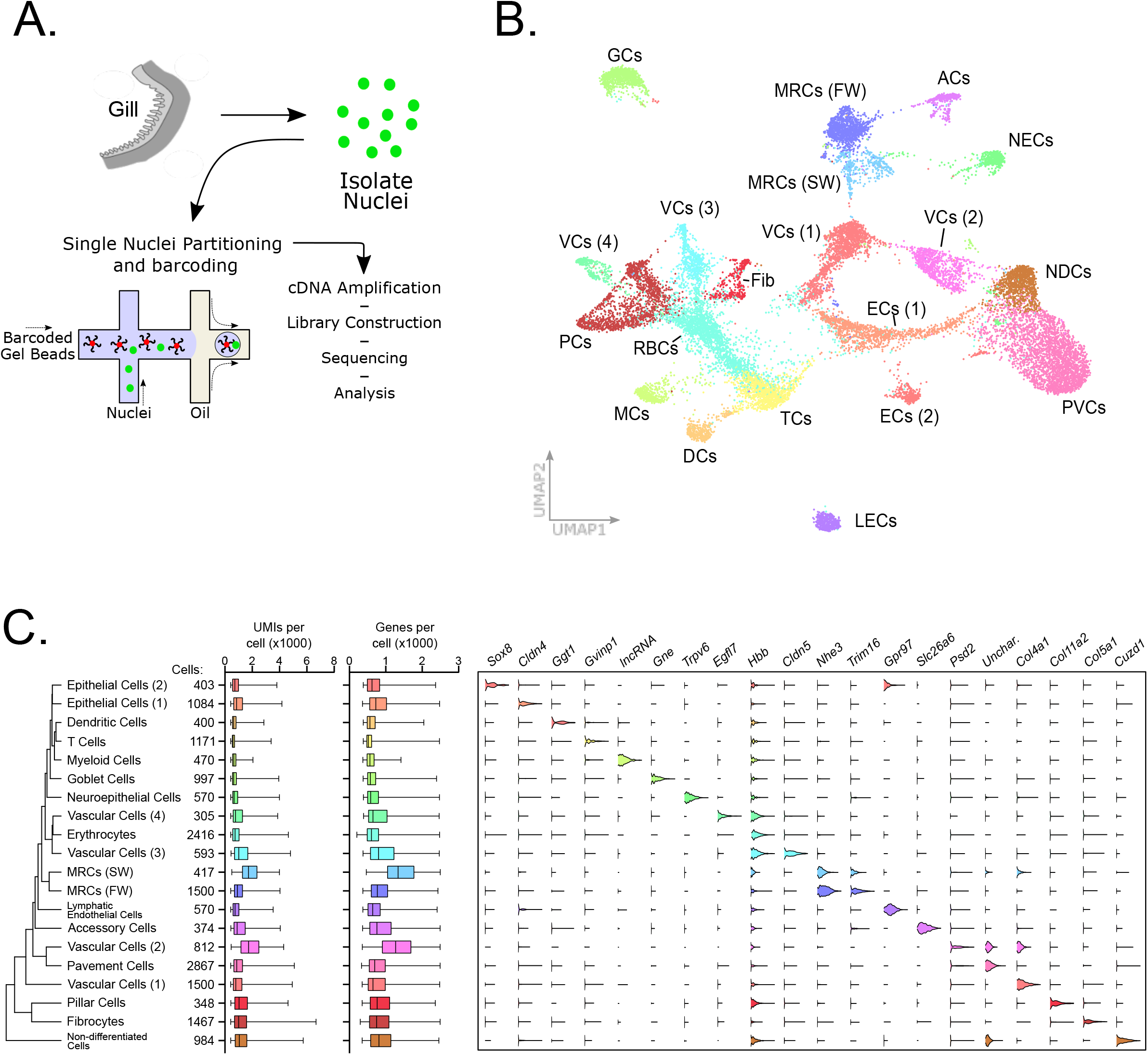
Single nuclei RNAseq analysis of Atlantic salmon gill tissue. A) Gill tissue processing. B) UMAP plot of pooled cell data from 18844 cells representing eight samples from four collection states. The plot indicates 20 separate cell clusters. C) Expression of marker genes in 20 cell clusters. From left to right: hierarchical relatedness of difference cell clusters; total cells in each cluster; UMI number in each cell cluster; gene features in each cell cluster; violin plots showing expression pattern of marker genes for each cluster. Abbreviations: ACs – accessory cells, DCs – dendritic cells, ECs – epithelial cells, fib – fibrocytes, GCs – goblet cells, LECs – lymphatic endothelial cells, MCs – myeloid cells, MRC – mitochondrion-rich cells, NDCs – non-differentiated cells, PVCs, pavement cells, RBCs – red blood cells (erythrocytes), TCs – T cells, VCs – vascular cells.

Lists of co-expressed marker genes define individual clusters. We categorized individual clusters using gene ontology analysis of marker gene lists and unique expression of known marker genes. This approach allowed us to infer cell types including fresh-and seawater MRCs, ACs, neuroendothelial cells, goblet cells, non-differentiated cells, pillar cells, lymphatic endothelial cells and several types of blood cell. We also identified a novel population of fibrocyte-like cells, and several types of vascular-and endothelial-like cells that partitioned across several clusters, together suggesting greater complexity in gill cytology that previously appreciated (Figure 1B).

We then defined expression signatures for each cell cluster. Our analysis re-identified several known marker genes, but also identifying several novel cell-type markers (Figure 1C). For example, the accessory cell signature included highly restricted expression of Slc26a6, an apical membrane Cl^−^/HCO_3_ exchanger, associated with gill function but until now misattributed to expression within MRCs ^25,26^. We were interested to note that the markers defining the erythrocyte population, including beta-globin, were expressed widely among all cell types. It is unclear exactly what role extra-erythroid haemoglobin plays in the gill, however, mammalian studies suggest that haemoglobin, in addition to its oxygen carrying capacity, may play an antimicrobial role ^27^. As a major mucosal immune barrier, this capacity may be pertinent to the gill ^28^.

### Major changes in cell composition during smoltification

To understand how gene expression and cellular complexity changes within the gill during smoltification and seawater transfer we compared the snRNAseq profiles at different developmental points (Fig 2A, for confirmation of smolt status see^29^). The abundance of six nuclei clusters changed dramatically (>3 fold change in percentage abundance) during smoltification (Figure 2B). SW MRCs increased in proportion steadily from T1-T4, consistent with previous descriptions of Atlantic salmon gill physiology. We also observed a marked increase in vascular cell number, with the major differences occurring between T2 and T3, suggesting that this vascular cell cluster proliferate in line with growth rates (Figure 2C). Interestingly, four immune-related nuclei clusters representing T cells, myeloid cells, dendritic cells and lymphatic endothelial cells fell dramatically during smoltification (Figure 2D). Changes in cell abundance occurred with a similar profile in all immune-associated cell clusters, with consistent decline observed between T1-T3. In contrast, 24h SW transfer does not appear to affect immune-cell abundance directly. These results highlight the complex and dynamic changes in cellular composition that occur in the gill during smoltification.

**Figure 2.**
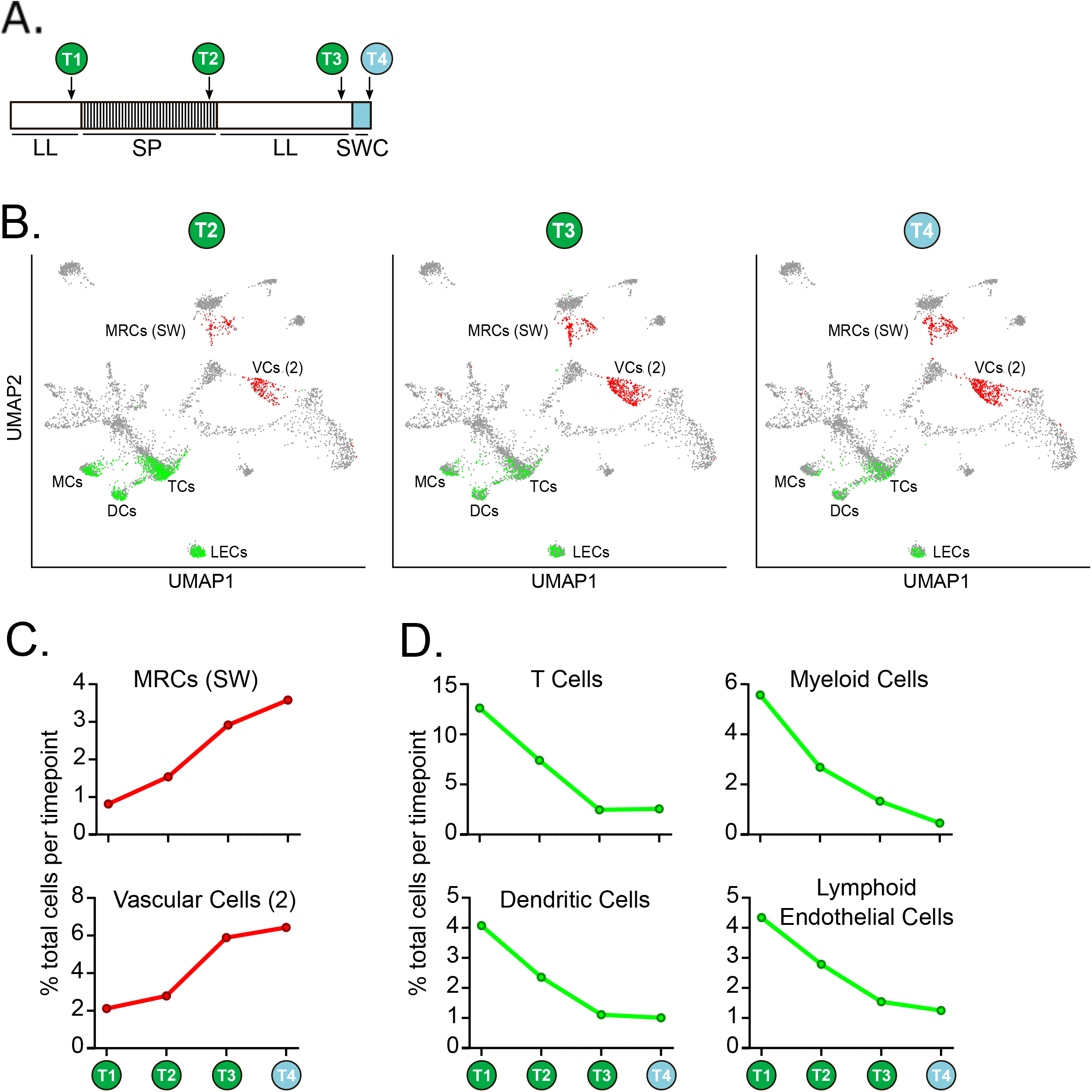
Comparative abundance of cell clusters at different sampling points. A) Experimental design. Fish were kept in constant light (LL) from hatching then transferred to short photoperiod (SP; 8L:16D) for 8 weeks before being returned to constant light (LL) for 8 weeks. Finally the fish were transferred to sea water for 24h. Sample points are indicated T1-T4. B) Subset of cell clusters from T2, T3 and T4 (green and red dots) overlaid on T1 cells (grey dots). C) Increasing abundance of sea-water mitochondrion-rich cells (MRCs SW) and vascular cells (VC 3) during smoltification C) Decreasing abundance of leukocytes and immune-associated cells during smoltification.

### Cell cluster-specific expression of smoltification-associated factors

Next, we wanted to identify cluster types where smoltification drives changes in local gene regulation. For statistical power, we cross-referenced our snRNAseq analysis with whole gill RNAseq analysis of T1-T3, identifying 9746 genes differentially regulated by smoltification (FDR <0.01). Pearson clustering of these genes resolved five major clusters that were associated with immune response, structural morphogenesis, autophagy, catabolism and mitochondrial respiration (Figure 3A). Within our analysis we identified a number of “classical” smoltification-related genes. As expected, CFTR was highest under constant light (LL), and was highly localized in expression to MRCs (Figure 3B). We also identified the reciprocal regulation of sodium-potassium ATPase subunits, specifically, suppression of NKAa1a and increase in NKAa1b ^12^. Inspection of cellular localization within our snRNAseq dataset showed that expression of these genes were, as anticipated, highest within the MRCs and ACs (Figure 3B).

**Figure 3.**
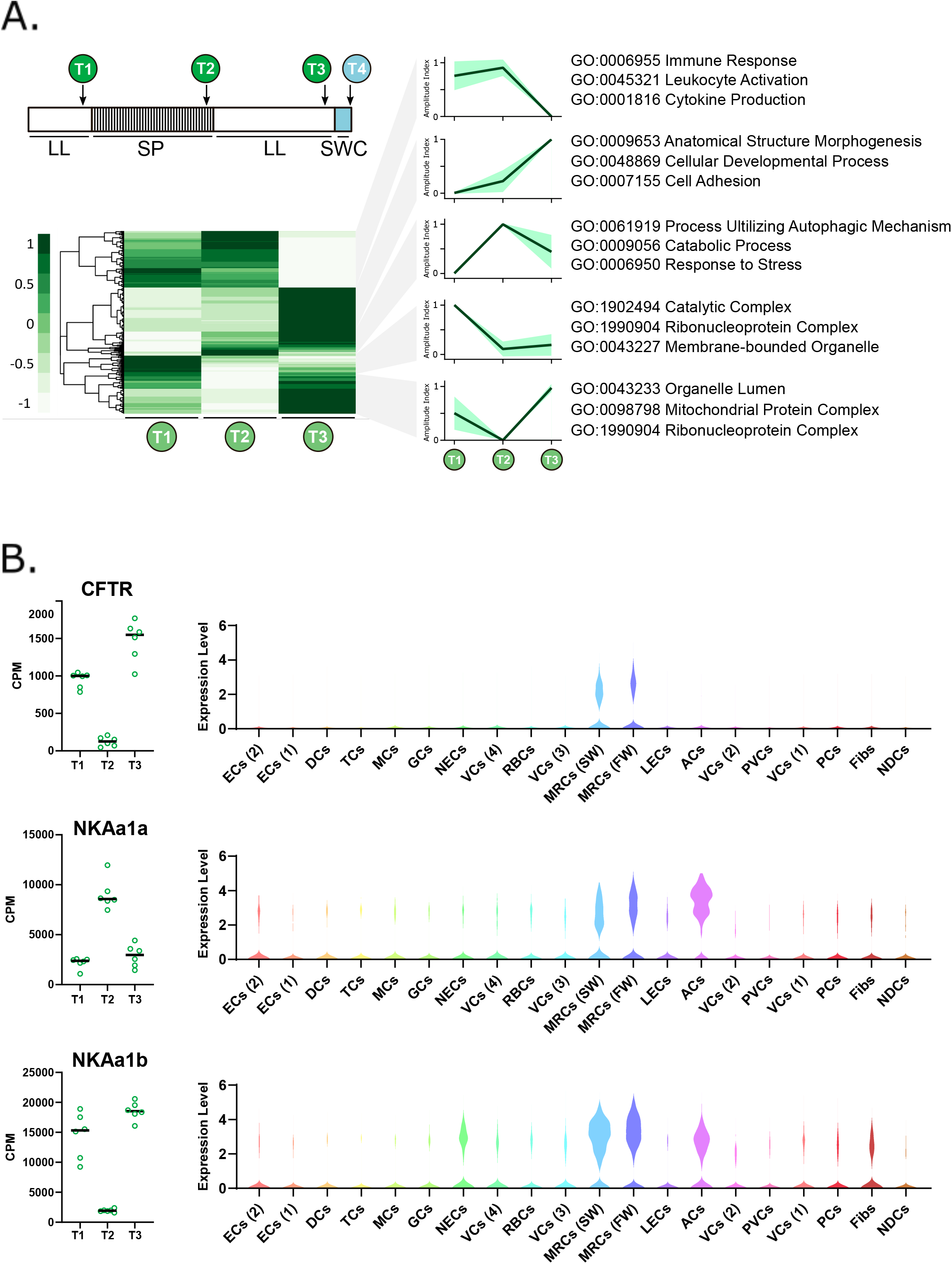
Photoperiodic changes in gill gene expression and localized cell cluster expression. A) Heat map representing 9746 genes differentially regulated (FDR <0.01) from T1-T3. Regulatory patterns for 5 major cluster are shown as amplitude index and 95% confidence limits. Major gene ontology terms for each cluster are shown. B) RNAseq data for “classical” smoltification-related genes and violin plots showing their cluster specific expression.

Our previous work identified genes whose expression are predicated on exposure to several weeks of short-photoperiod exposure^29^. In Atlantic salmon, these “winter-dependent” genes are analogous to vernalization dependent genes in *Arabidopsis*^30^, where a dosage of exposure to a winter-like stimulus (in *Arabidopsis*, cold; in Atlantic salmon, short photoperiod) controls the presentation of a seasonal phenotype under summer-like stimulus (in *Arabidopsis*, warmth and long days; in Atlantic salmon, long photoperiod). Winter-dependent genes are therefore intrinsically linked to unidirectional smolt development, and may play a mechanistic role in pre-adaptation of the gill for seawater migration. Surprisingly, canonical markers of smolt status, including the reciprocal expression of NKA subunits, are not winter dependent, meaning that their expression is passive to photoperiod rather than life history progression^29^.

Using our RNAseq dataset we identified novel, winter dependent genes. Next, we mined our snRNAseq dataset to identify the cell clusters that express these factors (Figure S1). Of particular interest was Cuzd1, a gene associated with carcinogenesis, whose expression was tightly localized to non-differentiated cells ^31^. We also identified Rhag, an ammonium transporter thought to be erythrocyte specific in mammals, but here predominantly expressed in the vascular cell (VC 3) cluster ^32^; and Hg2a (CD74) a ubiquitously expression multi-functional protein linked to immune defense ^33^. Taken together our data show that smoltification engages all gill cells in diverse regulatory phenotypes.

### Cell cluster-specific expression of seawater transfer-associated factors

Smoltification manifests when the Atlantic salmon smolts migrate downstream and arrive in the marine environment, thereby committing to an oceanic life ^1^. To gain insight into this critical step we compared RNA profiles of gill samples between smolts in freshwater and 24h in seawater using whole gill RNAseq, identifying 144 induced and 107 suppressed genes (FDR<0.01). Gene ontologies showed that the induced gene cohort was significantly associated with keratinization, whereas the suppressed gene cohort related to immune function (Figure 4A). We cross-referenced our whole gill RNAseq data against our snRNAseq data to isolate cell-type specificity of gene expression. These data highlight the restricted expression of key up-regulated genes (Figure 4B). For example, we localize the expression of an enzyme involved in both ionic and acid/base balance, carbonic anhydrase, to MRCs ^34,35^.

**Figure 4.**
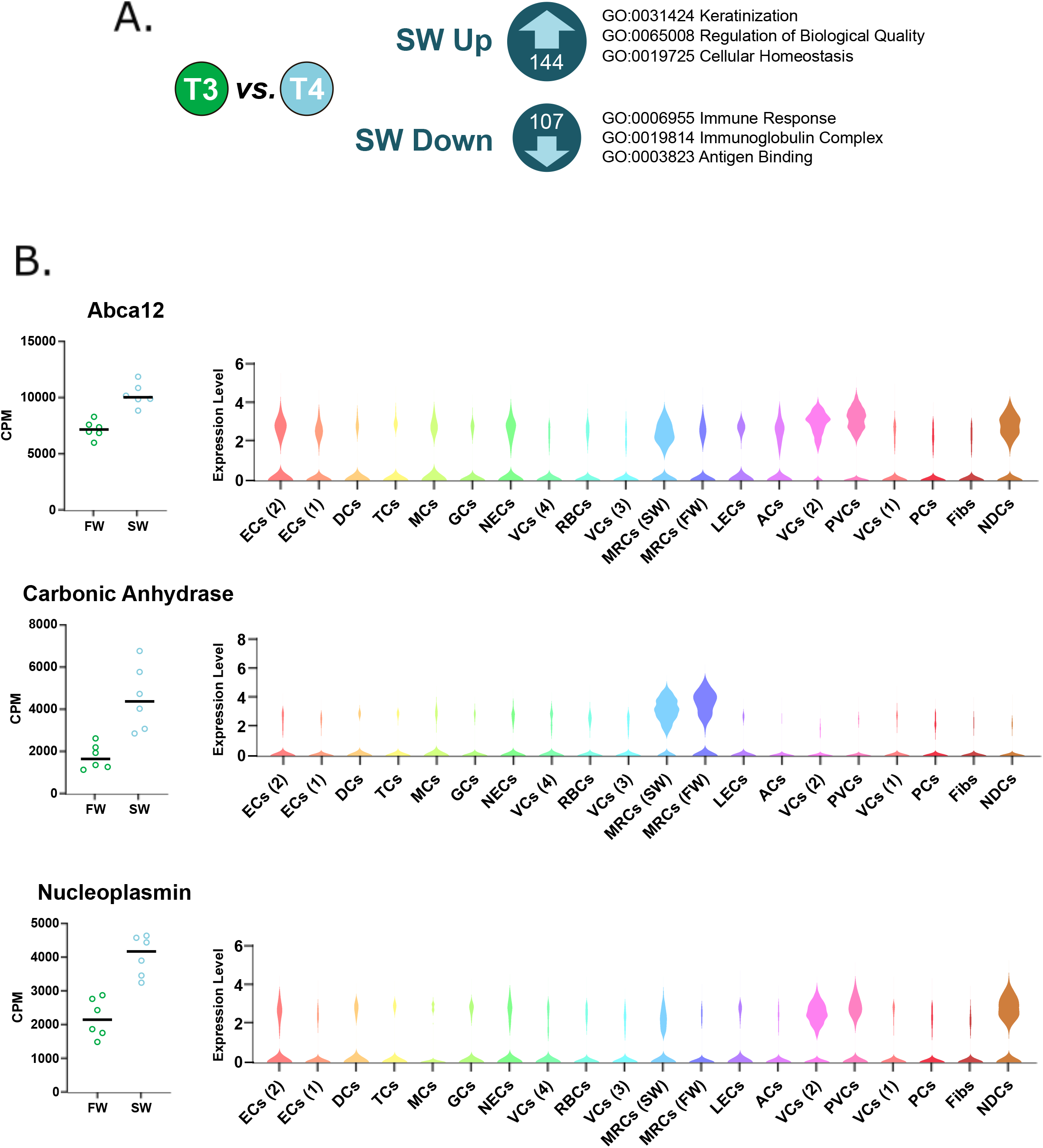
Sea-water transfer-associated changes in gill gene expression and localized cell cluster expression. A) Genes differentially regulated (FDR <0.01) by 24h seawater transfer. Major gene ontology terms for each cluster are shown. B) RNAseq data for sea-water transfer-related genes and violin plots showing their cluster specific expression.

We also show that ATP-binding cassette sub-family A member 12 (Abca12), a gene important in epidermal lipid barrier formation ^36^, is broadly expressed, but particularly concentrated in MRCs (SW), pavement, vascular, and non-differentiated cells. Interestingly, we show that a protein chaperone that helps regulate chromatin state, nucleoplasmin ^37,38^, is expressed specifically in non-differentiated, vascular and pavement cells groups, suggesting that these cell types undergo a change in chromatin status under seawater exposure.

## Conclusions

Our results bring insightful cellular resolution to the complexity of the Atlantic salmon gill and the compositional changes that occur during smoltification. Of particular interest was the suppression of immune cell types, which correlates with reduction in immune-related genes and suppression of immune function during smoltification ^39–41^. These data are a puzzle. The marine environment is awash with parasites, bacteria and viruses to which the salmon is potentially vulnerable, so loss of immune function would make little sense. Future work should focus on why and how the immune system is affected in aquaculture. Conceivably these data point towards an adaptive immunological reprogramming that helps to avoid immune shock when the salmon transition between the distinctive pathogen complements of fresh-and seawater habitats ^42,43^. Alternatively, artificial smolt production may drive abnormal immunosuppression. The constant light routinely used to stimulate smolts would profoundly undermine the immune defenses of mammals *via* disruption of the circadian clock ^44^.

Our data also shows that smoltification-driven transcriptional regulation occurs not only in MRCs and ACs, but also in other distinctive cell types including pavement cells, vascular cells and non-differentiated cells. We anticipate that novel gene function within the context of cell function will be a priority for future investigation, and will be assisted by the novel suite of marker genes which we present here.

## Material and Methods

### Animal welfare statement

The Atlantic salmon smoltification experiment was conducted as part of the routine, smolt production at Kårvik havbruksstasjonen, approved by the Norwegian Animal Research Authority (NARA) for the maintenance of stock animals for experiments on salmonids. This is in accordance with Norwegian and European legislation on animal research.

### Experimental Design

Atlantic salmon (*Salmo salar*, Aquagene commercial stain) were raised from hatching in freshwater, under continuous light (LL, > 200 lux at water surface) at ambient temperature (∼10°C). Juvenile salmon were housed in 500 L circular tanks and fed continuously with pelleted salmon feed (Skretting, Stavanger, Norway). At seven months of age parr (mean weight 49.5g) were sampled for T1 (experiment start). Two days later remaining parr were equally distributed between two 100L circular tanks, and over the next seven days the photoperiod was incrementally reduced to a short photoperiod (SP, 8h light:16h darkness). T2 sampling occurred on experimental day 53 (44 days on SP), remaining parr were transferred back to LL on experimental day 60. T3 sampling occurred on experimental day 110 (50 days after return to LL), then a sub-cohort of fish were netted out and transferred to full strength seawater for 24h before the final T4 collection.

### RNAseq Analysis

Gill samples were collected, RNA extracted and libraries prepared, sequenced and mapped as Iversen et al (2020). Raw counts were analysed using EdgeR (ver. 3.30.0), using R (ver. 4.0.2) and RStudio (ver. 1.1.456). An ANOVA-like test was used to identify differential expressed genes between T1-T3 samples. Clustering analysis was performed using Pearson correlation, and heapmaps rendered using the R package pheatmap. An exact test was performed to identify differential expressed genes between T3 and T4.

### Single nuclei RNAseq Analysis

Gills for single nuclei analysis were collected on dry ice and stored at −80°C. Duplicate samples were processed for T1-T4. Nuclei were released by detergent mechanical lysis, then samples were homogenized (30s) and nuclei isolated by sucrose gradient ^45^. Libraries were created using Chromium Single Cell 3’ GEM, Library & Gel Bead Kit v3 (10x technologies) by University of Manchester genomic technology core facility (UK). Raw data was converted to counts per cell using Cell Ranger (10x Technologies, ver. 3.1.0) and processed using NCBI annotations. The R package Seurat (ver. 3.1.5) was used to perform an integrated analysis using all snRNAseq data ^24^, further details in results and discussion. Raw data will be available following peer-reviewed publication.

## Acknowledgements

The authors thank all of the animal staff at Kårvik havbruksstasjonen for their expert care of the research animals, and the University of Manchester Genomics Technology core facility (UK) for performing chromium 10x library preparation for snRNAseq. ACW is supported by the Tromsø forskningsstiftelse (TFS) grant awarded to DGH (TFS2016DH). The Sentinel North Transdisciplinary Research Program Université Laval and UiT awarded to DGH supports this work. SHW is supported a grant from the Tromsø forskningsstiftelse (TFS) starter grant TFS2016SW. Experimental costs were covered by HFSP grant “Evolution of seasonal timers” RGP0030 /2015 awarded to ASIL and DGH.

## Author Contributions

Conceptualization, A.C.W., Y.M., E.H.J., A.S.I.L and D.G.H; Resources, A.C.W, Y.M and M.I.; Investigation, A.C.W., Y.M., M.I. E.H.J and D.G.H; Formal Analysis, A.C.W, Y.M., L.M.I., T.N and S.R.S; Visualization, A.C.W and S.H.W; Writing – Original Draft, A.C.W and S.H.W; Writing – Review & Editing, All; Supervision, A.S.I.L and D.G.H; Project Administration, A.S.I.L and D.G.H; Funding Acquisition, A.S.I.L and D.G.H.

---

**Supplemental Figure 1**. RNAseq data for winter-dependent genes and violin plots showing their cluster specific expression.

## Notes

### Competing Interest Statement

The authors have declared no competing interest.

