## Supplementary figures and images for "A single nuclei transcriptomic analysis of the Atlantic salmon gill through smoltification and seawater transfer"

### Supplemental Figure 1

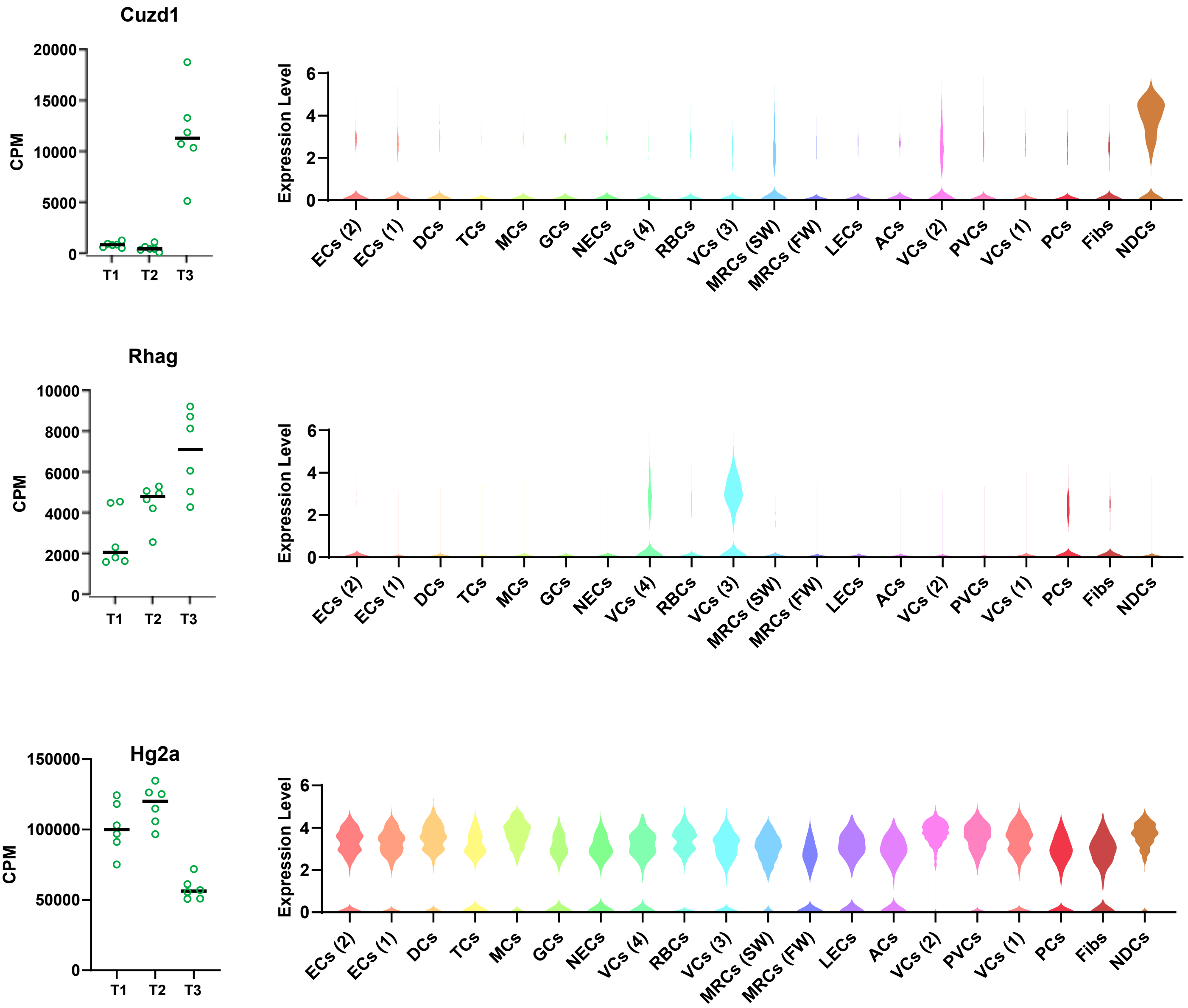
